# Systematic Analysis of the Glucose-PTS in *Streptococcus sanguinis* Highlighted its Importance in Central Metabolism and Bacterial Fitness

**DOI:** 10.1101/2024.07.25.605150

**Authors:** Zachary A. Taylor, Danniel N. Pham, Lin Zeng

**Affiliations:** Department of Oral Biology, University of Florida College of Dentistry, Gainesville, FL 32610, USA

## Abstract

Previous work reported that deletion of the Enzyme IIAB subunits (EIIAB^Man^, *manL*) of the glucose phosphotransferase system (glucose-PTS, *manLMNO*) in *Streptococcus sanguinis* impacted carbon catabolite repression (CCR) and bacterial fitness. Here, a single nucleotide polymorphism in ManN, ManNA91E, produced the unusual phenotype of increased excretion of organic acids and H_2_O_2_, yet elevated PTS activities. To characterize the contributions of each component of the glucose-PTS to bacterial fitness, we performed genetic analyses by deleting from *S. sanguinis* SK36 the entire operon and each EII^Man^ subunit individually; and genes encoding the catabolite control protein CcpA (Δ*ccpA*) and the redox regulator Rex (Δ*rex*) for comparison. Deletion of each subunit incurred a growth defect on glucose partly due to elevated excretion of H_2_O_2_; when supplemented with catalase this defect was rescued, instead resulting in a significantly higher yield than the parent. All glucose-PTS deletion mutants presented an increased antagonism against the oral pathobiont *Streptococcus mutans*, a phenotype absent in Δ*ccpA* despite increased H_2_O_2_ output. A shift in the pyruvate node towards mixed acid fermentation and increased arginine deiminase activity enhanced pH homeostasis in glucose-PTS mutants, but not Δ*ccpA*. Despite the purported ability of Rex to regulate central carbon metabolism, deletion of *rex* had no significant impact on most of the phenotypes discussed here. These findings place glucose-PTS in the pivotal position of controlling central carbon flux in streptococci, with critical outcomes impacting acidogenicity, aciduricity, pH homeostasis, and antagonism, highlighting its potential as a therapeutic target for treating diseases with a dysbiotic microbiome.

**Importance:** Management of carbohydrate metabolism and environmental stress is key to the survival of oral commensal species such as *S. sanguinis*. Antagonism of oral pathobionts and modulation of the environmental pH and oxidative potential by commensals are crucial to the maintenance of microbial homeostasis and prevention of oral diseases including dental caries. It is therefore vital to understand how these species regulate sugar fermentation, production of acids and ammonia, and stress management in an environment known for a feast-and-famine cycle of carbohydrates and similar fluctuations in pH and oxygen tension. Here, we detail that genetic alterations of the glucose-PTS transporter in *S. sanguinis* can significantly affect the regulation of factors required for bacterial fitness and homeostatic ability independent of known catabolic regulators, it is then discussed how these changes may impact the survival of streptococcal species and affect caries onset.

## Introduction

Dental caries is the most prevalent chronic disease globally (1). As it is considered a polymicrobial disease characteristic of an over-abundance of multiple highly acidogenic and aciduric microorganisms (2–7), it is necessary to view caries in the greater context of the oral microbiota (8). While oral biofilms are incredibly diverse (2, 9), *Streptococcus* species are of particular importance. Their relative abundance, ability to ferment a variety of carbohydrates and produce organic acids, and the status of many as early colonizers (10, 11), are central to balancing the oral environment between a healthy and diseased state. One of the main etiological agents of caries is the pathobiont *Streptococcus mutans*, known for its extraordinary acidogenicity and aciduricity that drives microbial dysbiosis and contributing to caries initiation and progression (4, 12). On the other hand, prevalence of the oral commensals such as *Streptococcus sanguinis, Streptococcus gordonii,* and certain members of the mitis group streptococci is associated with enhanced health due to their ability to secrete H_2_O_2_ that inhibits pathobionts such as *S. mutans*, and ammonia that favors commensals by raising the environmental pH (13, 14). Collectively, these characteristics of the commensal streptococci aid in maintaining microbial diversity and contributing to pH homeostasis associated with a healthy oral microbiome (7, 15).

In oral streptococci, these competitive factors are largely regulated by carbohydrate availability, a phenomenon known as carbon catabolite repression (CCR) (16). It has been established that catabolite control protein A (CcpA) is the primary transcription regulator of CCR in low-GC, Gram-positive bacteria such as *Bacillus subtilis*, wherein it works in association with a heat-stable phosphocarrier protein HPr in response to metabolic intermediates such as fructose-1,6-bisphosphate (FBP) (17). Previous work has established the broad transcriptional control CcpA incurs on a multitude of cellular functions including energy metabolism, extracellular polysaccharide (EPS) production, H_2_O_2_ excretion, pH homeostasis, and virulence expression (18–25). However, research in *S. mutan*s and other streptococcal pathogens has suggested that the CcpA-central model of transcriptional control represents an incomplete picture of CCR in streptococcal pathophysiology (23, 26–29). For example, the phosphoenolpyruvate::sugar phosphotransferase system (PTS) of *S. mutans* can work in concert with CcpA, as well as independently, to exert a regulatory function for a multitude of catabolic genes (23, 30, 31). We recently discovered that this regulatory role was translatable to *S. sanguinis*, with the identification of spontaneous truncation mutants of the enzyme II (EIIAB^Man^, *manL*) of the glucose-PTS which presented altered cellular processes and fitness phenotypes beyond sugar transport (32). In fact, mutants lacking an intact *manL* had increased expression of pyruvate oxidase (*spxB*) (33) leading to increased H_2_O_2_ excretion comparable to a *ccpA* mutant. This resulted in increased antagonism against *S. mutans* by *manL* mutants, a phenotype absent in the *ccpA* mutant (34). Further RNAseq analysis in a *manL* mutant revealed altered expression in a 311-gene regulon encompassing secondary carbohydrate metabolism, pyruvate metabolism, alkali production, membrane biosynthesis, and attachment, with limited overlap to systems affected by the deletion of *ccpA* (32).

A multitude of metabolic processes depend on the balanced redox state of the cell, which in Gram-positive bacteria is partly regulated by the redox regulator Rex (35–38). Rex has been shown to regulate several operons that help to maintain NAD^+^/NADH balance in lactic acid bacteria, many of which deal with the later steps of glycolysis (36), including the reduction of pyruvate to lactate by lactate dehydrogenase that regenerates NAD^+^ from NADH (39). While the Rex regulon has been delineated in streptococci bioinformatically, its actual functions are not well characterized, since a deletion of the *rex* gene in *S. mutans* did not affect LDH expression as predicted (35). As *manL* deletion also affected several genes classified for the Rex operon (32), the relationship between these two systems requires clarification.

To understand the underlying mechanisms and significance of PTS-dependent regulation in streptococcal physiology, we conducted a systematic genetic analysis, first by passaging *S. sanguinis* SK36 under the condition from which the *manL* truncation was initially identified, then conducting mutagenesis that targeted the individual subunits of the glucose-PTS EII (*manLMN*), an ORF of unknown function (*manO*), as well as major metabolic regulators including CcpA, HPr, and Rex. Results obtained while characterizing these mutants highlighted the central role of glucose-PTS in streptococcal fitness by regulating energy metabolism, pH homeostasis, and H_2_O_2_-mediated antagonism.

## Results

### An SNP mutant of *manN* exhibited phenotypes indicative of CCR relief despite increased PTS activity

To understand the emergence of previously identified spontaneous mutations in the glucose-PTS (*manL*), *S. sanguinis* wildtype (WT) lab strain SK36 was passaged in BHI for 15 days, with media refreshment once every 24 hours. The resulting populations were assayed for H_2_O_2_ excretion utilizing a Prussian Blue (PB) plate assay. Compared to the starting stock, the passaged populations displayed a significant increase in PB zone when growing on glucose, but not on lactose, a sugar not transported by the glucose-PTS (Fig 1). Whole genome sequencing (WGS, Table S1) revealed several single nucleotide polymorphisms (SNP), one of which resided in the *manN* gene (EIID) of the glucose-PTS, a switch of the 91^st^ alanine to glutamic acid, and was present in approximately 95% of one of the sequenced populations (RD15-2). Other alterations included missense mutations in a hypothetical protein (SSA_0815) and a pseudogene (SSA_1885) (Table S1).

**Fig 1.**
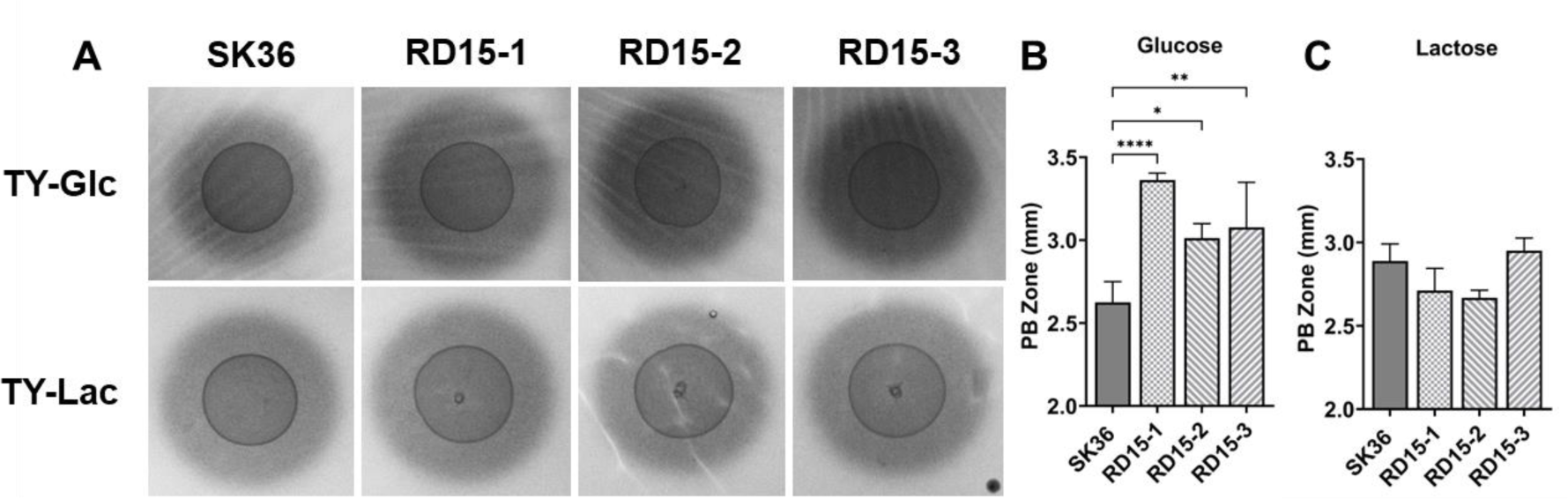
H_2_O_2_ production of SK36 and passaged populations. (A) Cultures of SK36 wild type and three populations of SK36 that had been passaged for 15 days (RD15-1 to 3) were each dropped onto TY-Prussian Blue (PB) agar plates supplemented with either 20 mM glucose (B) or lactose (C), and incubated for 24 hours in a 5% CO_2_ environment. All images were photographed under the same settings. Each PB zone was measured from the edge of the bacterial colony to the edge of the PB precipitation at four locations using ImageJ software. Asterisks represent statistical significance compared to the wild type according to one-way ANOVA (*, *P* <0.05; **, *P* <0.01; ***, *P* <0.001; ****, *P* <0.0001).

The *manNA91E* mutation was reconstituted in the WT SK36 background via site-directed mutagenesis and the resultant strains were labeled ManNA91E and subjected to WGS. Several SNPs were identified in each of the two isolates we sequenced, although none was shared other than *manNA91E* (see Table S2 for WGS data). Both isolates showed comparable phenotypes in the following studies. First, RT-qPCR was performed to assess the expression of genes required for production of organic acids, ammonia, and H_2_O_2_. Heterolactic fermentation seemed to be favored by an increased expression of acetate kinase (*ackA*) and pyruvate-formate lyase (*pfl*), albeit little change in expression by lactate dehydrogenase (*ldh*) (Fig 2A). Notably, *arcA* of the arginine deiminase system (ADS) responsible for arginine deimination and ammonia secretion was upregulated, as was pyruvate oxidase (*spxB*) responsible for the majority of H_2_O_2_ production while growing on glucose. Both genes are known to be negatively regulated by CcpA in oral streptococci (18, 19, 40, 41).

**Fig 2.**
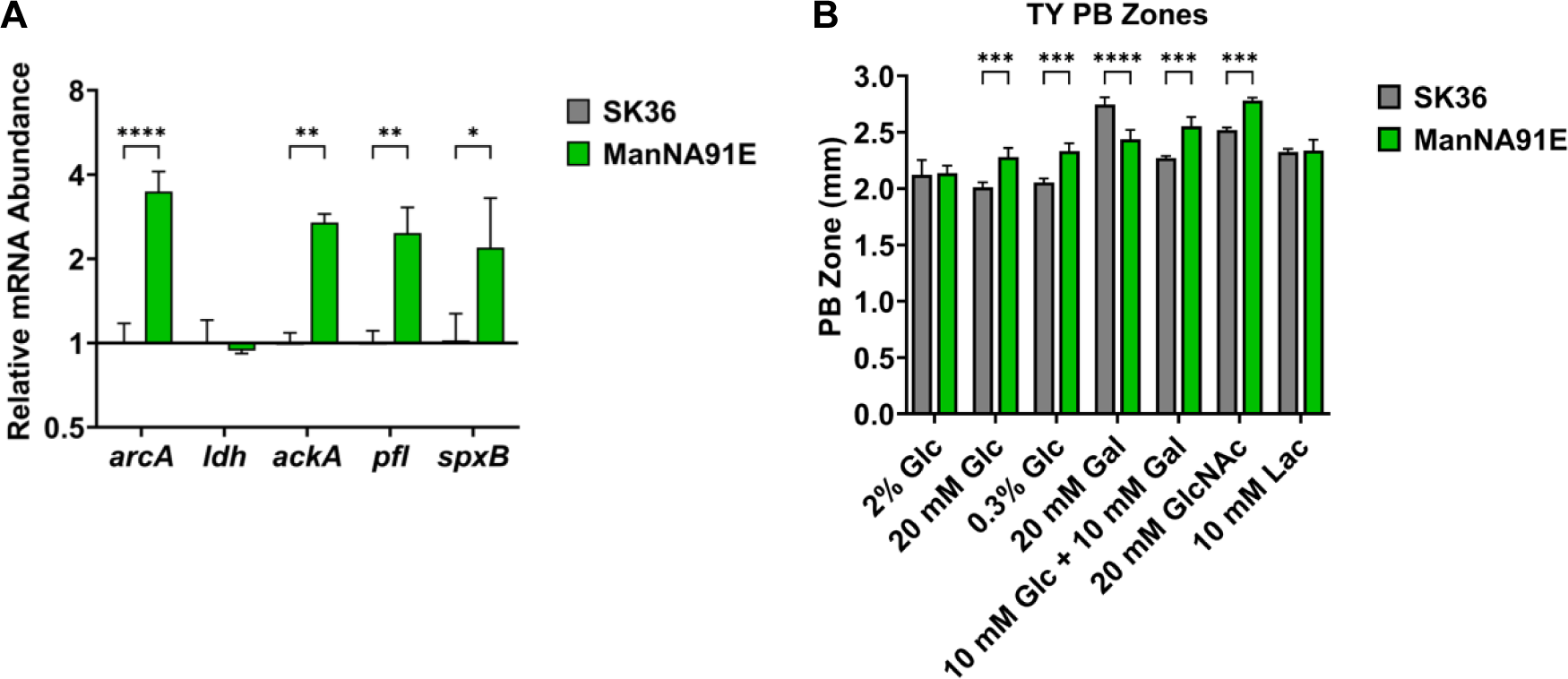
Transcription of metabolic genes (A) and H_2_O_2_ excretion (B) by SK36 and ManNA91E. To measure abundance of mRNA levels of metabolic genes, SK36 and its mutant derivative were grown to exponential phase in tryptone-yeast medium supplemented with 20 mM of glucose. RNA was extracted from cells and relative abundance was calculated relative to an internal control (*gyrA*) (A). For quantification of H_2_O_2_ excretion, 10 µl of cells were spotted onto TY agar plates supplemented with various sugars and incubated for 24 hours in a 5% CO_2_ environment. Each PB zone was measured from the edge of the bacterial colony to the edge of the PB precipitation at four locations using ImageJ software. Asterisks represent statistical significance compared to the wild type according to two-way ANOVA (*, *P* <0.05; **, *P* <0.01; ***, *P* <0.001; ****, *P* <0.0001).

Phenotypic characterization was conducted to assess the physiological impact of *manNA91E* mutation. Like the passaged populations (Fig 1), strain ManNA91E displayed, compared to SK36, an increased excretion of H_2_O_2_ on PB plates containing glucose, as well as N-acetylglucosamine (GlcNAc) which is another substrate of streptococcal glucose-PTS, yet with no change when growing on lactose plates (Fig 2B). A reduction in PB zone by ManNA91E was noted on galactose plates. We then measured organic acids in supernatants of cultures grown in a tryptone-vitamin (TV) medium supplemented with glucose (TVG). When compared to SK36, metabolic acid excretion from ManNA91E was increased for heterolactic (acetate, formate), but not homolactic (lactate) end products (Fig 3ABC). The sum of all three major acids showed a notable increase in ManNA91E (40.0 ± 1.7 vs 37.1 ± 2.1 mM/OD_600_), yet an increase in the resting pH of the same culture was observed (Fig 3D). Taken together with the transcription analysis, carbon flux through the pyruvate node appeared to have shifted from a mainly homolactic strategy to a mixed acid fermentation strategy, with increased acid production being offset by an increased arginine deiminase activity that excreted pH-buffering ammonia. This type of metabolic shift is often considered an indication of alleviation in CcpA-mediated CCR and an outcome of decreased carbon flux (16, 19, 21).

**Fig 3.**
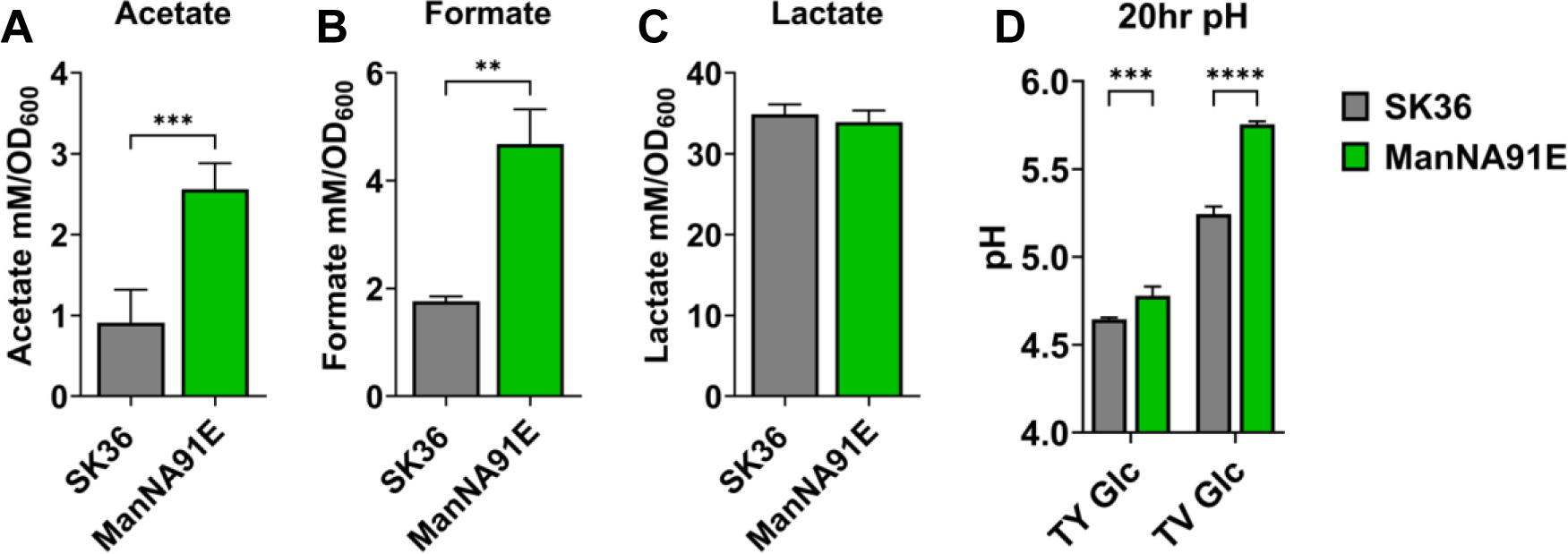
Metabolic acid excretion (A-C) and culture pH (D) of SK36 and ManNA91E. Cells were grown in TV (A-C) or TY (D) medium supplemented with 20 mM glucose to mid exponential phase for enzymatic assays (A-C) or for 20 hours for pH measurement (D). Asterisks represent statistical significance compared to the wild type according to Student’s *t*-test (A-C) or two-way ANOVA (D) (*, *P* <0.05; **, *P* <0.01; ***, *P* <0.001; ****, *P* <0.0001).

To corroborate this theory, ManNA91E was assayed for growth in Tryptone-Yeast extract (TY), TV, or a chemically-defined FMC medium, each supplemented with glucose as the sole carbon source. Other than a slight increase in lag on TY- and TV-based media, however, ManNA91E did not exhibit an altered growth phenotype indicative of a decreased carbon flux (Fig 4A-B), which was especially clear when assayed in FMC-glucose (Fig 4C). On the contrary, when tested in FMC containing 10 mM galactose, strain ManNA91E presented significant improvements in both growth rate and final yield compared to the wild type (Fig 4D). To confirm these growth phenotypes, PTS activities were measured using permeabilized cells. In support of the enhanced growth in FMC-galactose, *in vitro* phosphorylation of galactose by ManNA91E was significantly higher, with levels twice that of the wild type (Fig 4E). Similar results were obtained when other known substrates of the glucose-PTS, glucose, GlcN, and GlcNAc, were tested in the same assay. Finally, two isolates of ManNA91E were tested in studies described here, both showing comparable results (Fig S1-3).

**Fig 4.**
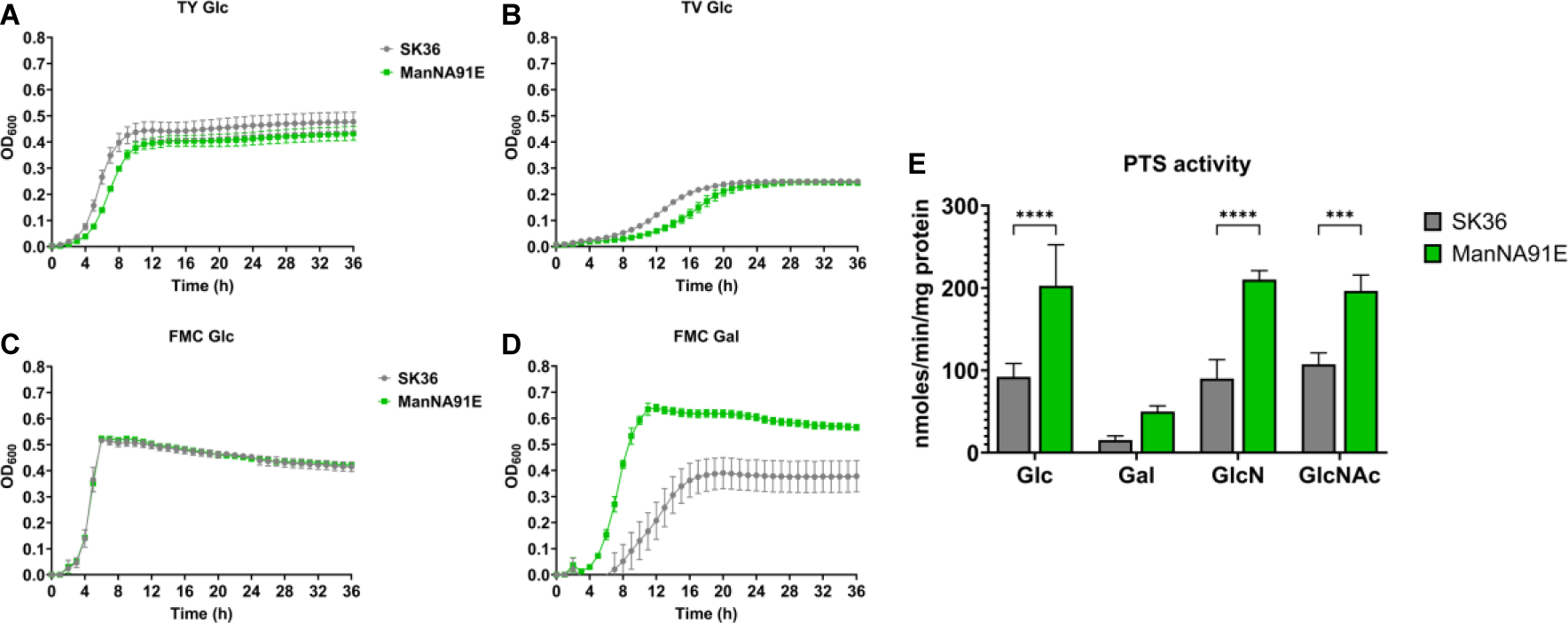
Growth curves (A-D) and PTS activities (E) of SK36 and ManNA91E. To measure growth, strains SK36 and ManNA91E were first cultured to mid-exponential phase in BHI and then diluted 1:100 into TY (A), TV (B), or FMC (C-D) medium supplemented with glucose (A-C) or galactose (D). To measure PTS activity, SK36 and ManNA91E cells were cultured in BHI medium, harvested from mid-exponential phase, and subjected to an *in vitro* sugar phosphorylation assay (E). Results are each an average of at least three biological replicates. Asterisks represent statistical significance compared to the wild type according to Student’s *t*-test (*, *P* <0.05; **, *P* <0.01; ***, *P* <0.001; ****, *P* <0.0001).

Considering the presence of several spontaneous SNPs identified in these isolates (Table S2), we took this one step further, by identifying the conserved, corresponding amino acid residue of ManNAla91 in the orthologous gene of SGO_1681 that encodes a *manN* homolog in a closely related, metabolically similar *S. gordonii* strain DL1 and constructed a DL1_manNA81E mutant. Remarkably, strain DL1_manNA81E showed similarly enhanced growth on galactose compared to DL1 and produced higher amounts of H_2_O_2_ on PB plates supported by glucose, GlcNAc, or a combination of glucose and galactose, but not on lactose alone (Fig S4).

Together, these findings with ManNA91E SNP led us to posit that the glucose-PTS is a significant factor in the regulation of central metabolism and bacterial fitness, as CcpA-mediated CCR is insufficient to explain these novel phenotypes (16, 32). To further test this hypothesis and overcome the limitation related to the presence of other spontaneous SNPs, deletion mutants of individual glucose-PTS subunits were genetically constructed and studied along with the mutant of *ccpA.* A deletion mutant of *rex* was also constructed given its suggested function in regulating metabolism in response to redox signals (38, 42).

### Glucose-PTS EII deletion mutants in *S. sanguinis* SK36 displayed altered growth rate and yield in multiple carbohydrates

To compare the growth phenotypes of SK36 mutants with a deficiency in various glucose-PTS subunits, strains Δ*manL* (EIIAB^Man^), Δ*manM* (EIIC^Man^), Δ*manN* (EIID^Man^), Δ*manO*, Δ*manLMNO* (EIIABCD^Man^/ManO), Δ*rex*, and Δ*ccpA* were grown in TY medium supplemented with glucose, galactose, glucosamine (GlcN), or GlcNAc, all of which are glucose-PTS substrates. In TY-glucose (Fig 5), each EII^Man^ mutant displayed a slower growth rate, i.e., an increase in doubling time (T*_d_*, Table S3) ranging from 8 to 42 minutes. Significantly, the final yield of these same EII^Man^ mutants each increased by about ΔA=0.3-0.4 OD_600_. The Δ*ccpA* strain shared this increase in yield with a ΔA=0.2 OD_600_ but showed a significantly longer lag phase. This reduction in growth rate by the EII^Man^ mutants was more drastic when galactose was substituted for glucose in TY medium (Fig S5A), leading to increases in T*_d_* values ranging from 100 to 130 minutes (Table S3). Similar to galactose, reduced growth rates were presented by these EII^Man^ mutants when tested in GlcN or GlcNAc, as did Δ*ccpA* (Fig S5B&C). Different from glucose though, deletion of each or all these EII^Man^ subunits resulted in reduced yields in each of these three sugars (Fig S5 and Table S3); Δ*ccpA* produced a reduced yield in galactose and especially in GlcN, but again had a significant lag phase increase in each sugar. Δ*manO* behaved in a manner close to the WT parent in glucose and galactose, however produced a slower growth rate yet a higher yield in both GlcN and GlcNAc; all EII^Man^ subunit mutants produced lower yields on either amino sugar. Of note, the Δ*rex* mutant lacked any discernible growth phenotype, mirroring the WT in all conditions tested.

**Fig 5.**
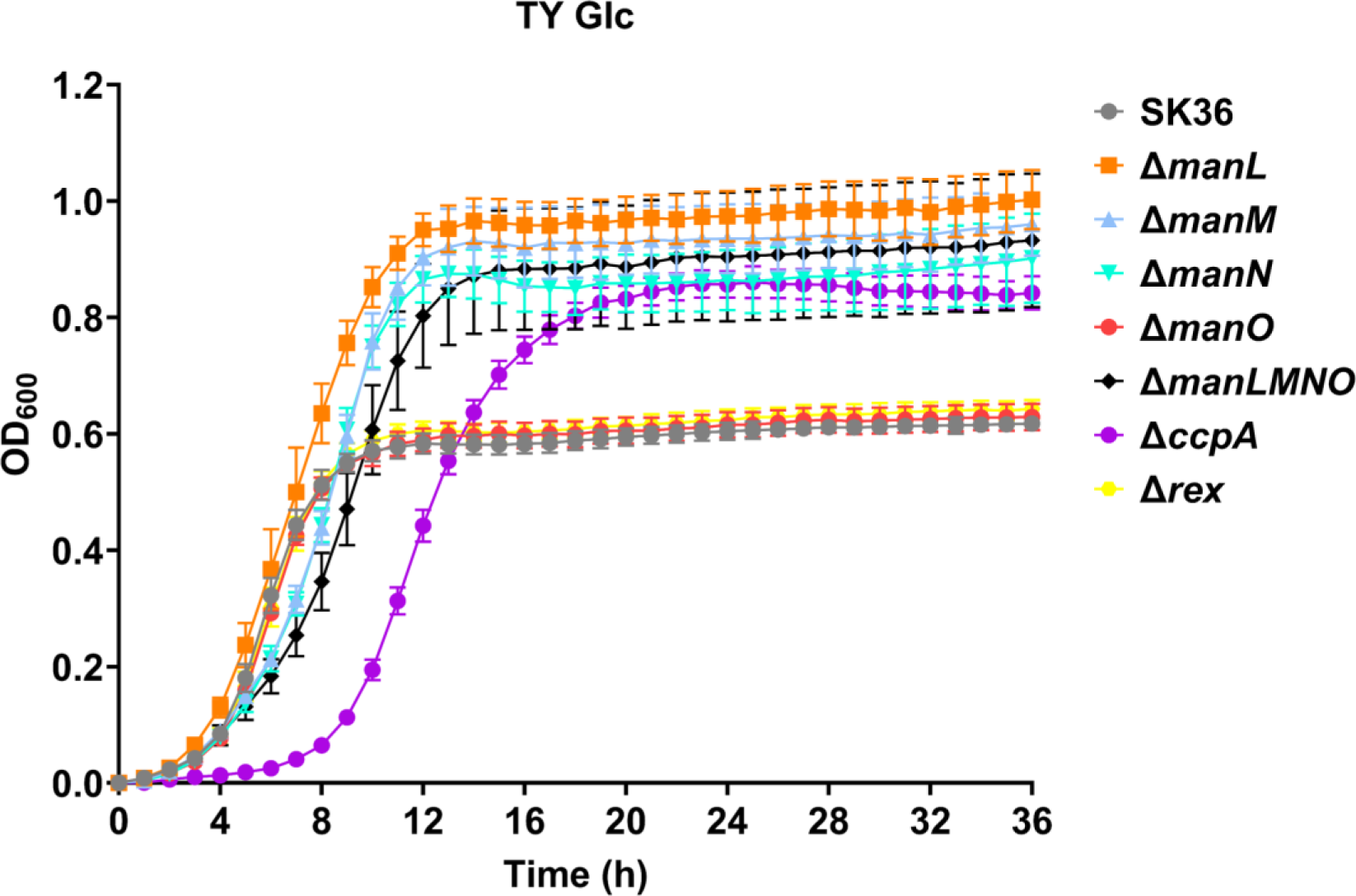
Growth curve of SK36 and various deletion mutants. Strains were first cultured to exponential phase in BHI and then diluted 1:100 into fresh TY medium containing 20 mM of glucose. Optical density at 600 nm (OD_600_) was monitored using a Bioscreen C over the course of 36 hours. Results are each an average of at least four biological replicates.

### Deletion of glucose-PTS EII subunits enhanced H_2_O_2_ excretion yet decreased eDNA release

The ability of *S. sanguinis* to antagonize pathobionts, e.g., *S. mutans,* in the oral cavity comes in large part through the secretion of H_2_O_2_, which is heavily regulated by carbon availability and CcpA activity (40). We measured the production of H_2_O_2_ by each deletion mutant of the EII^Man^ using both an enzymatic assay and the PB plate assay. There was a general increase in H_2_O_2_ levels in the supernatants of these mutants relative to the WT, with Δ*manM* showing the greatest increase and Δ*manO* the least (Fig 6A). Δ*ccpA* produced H_2_O_2_ at levels comparable to that of most PTS mutants, yet little change was observed in Δ*rex* (Fig 6A). This across-the-board increase in H_2_O_2_ production by these mutants was substantiated by an equally significant, though not all to the same degree, increase in the mRNA levels of the pyruvate oxidase (*spxB*) (Fig 6B), with Δ*ccpA* producing the highest level. The PB plate assay (Fig S6) largely confirmed the findings made in the liquid cultures.

**Fig 6.**
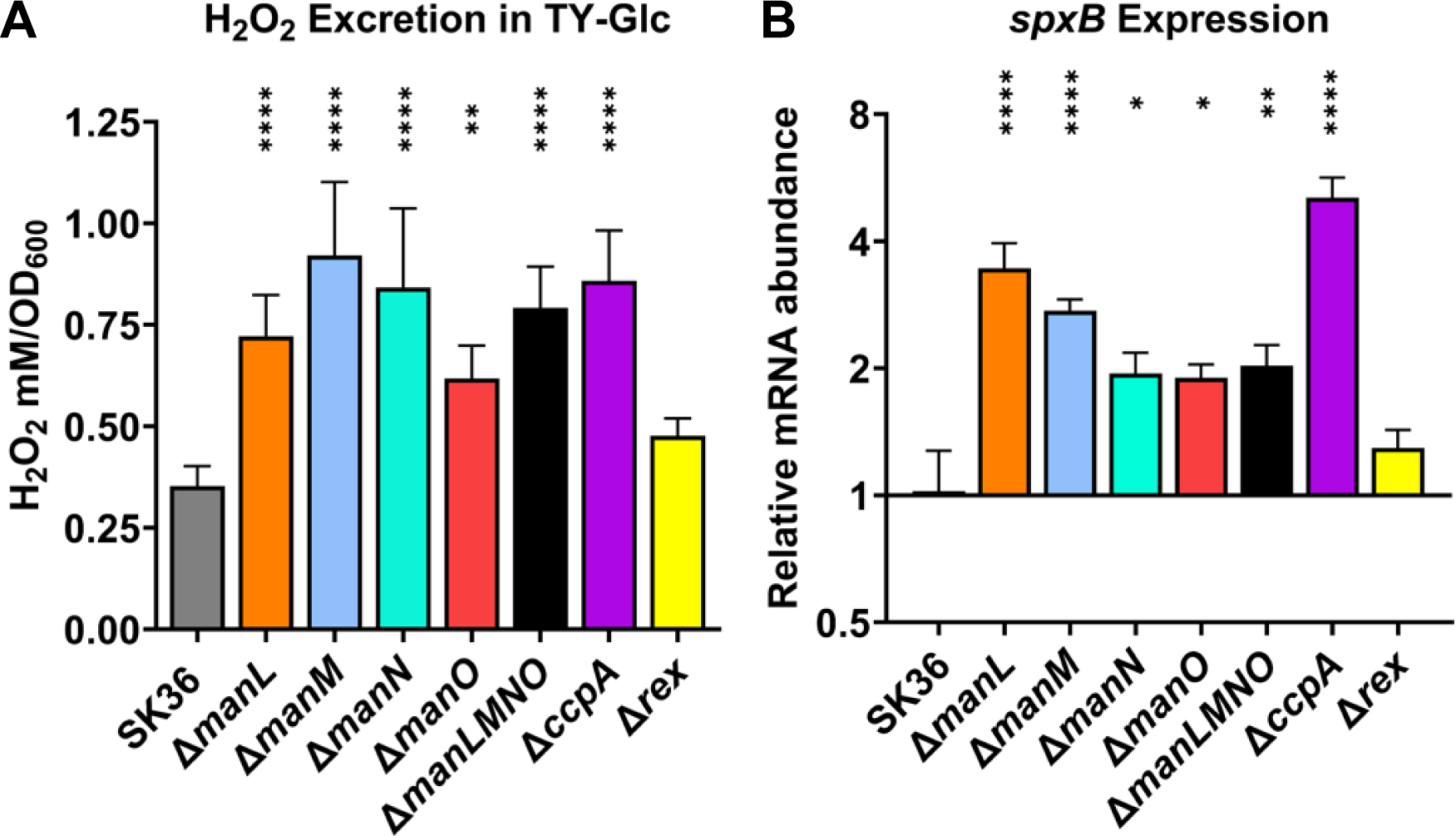
H_2_O_2_ excretion (A) and expression of pyruvate oxidase *spxB* (B) of SK36 and mutant derivatives. (A) strains were grown to early-exponential phase in TY-glucose, shaken at 250 RPM for 30 minutes in aerobic atmosphere before being tested using a colorimetric assay and a H_2_O_2_ standard curve. (B) cells from mid exponential phase were harvested for RNA extraction, followed by reverse transcription and qPCR using an internal control (*gyrA*). Results are the averages of three biological replicates, with error bars denoting standard deviations. Asterisks represent statistical significance compared to the wild type according to one-way ANOVA (*, *P* <0.05; **, *P* <0.01; ***, *P* <0.001; ****, *P* <0.0001).

Previous research has shown that elevated excretion of H_2_O_2_ does not always correlate with a greater ability to antagonize other bacteria (43). To test their capacity in interspecies antagonism, each mutant was spotted side by side with *S. mutans* strain UA159 on TY agar plates supplemented with glucose (TY-Glc) or lactose (TY-Lac), or on BHI agar plates. The resulting inhibition of UA159 by all glucose-PTS mutants (excluding Δ*manO*) was greater than the WT on TY-Glc and BHI plates, but not on TY-Lac plates (Fig 7 and Fig S7). Like the WT, Δ*rex* showed little activity in antagonizing UA159 on all three plates, so did Δ*ccpA* despite its excreting the greatest levels of H_2_O_2_, as was reported previously (34).

**Fig 7.**
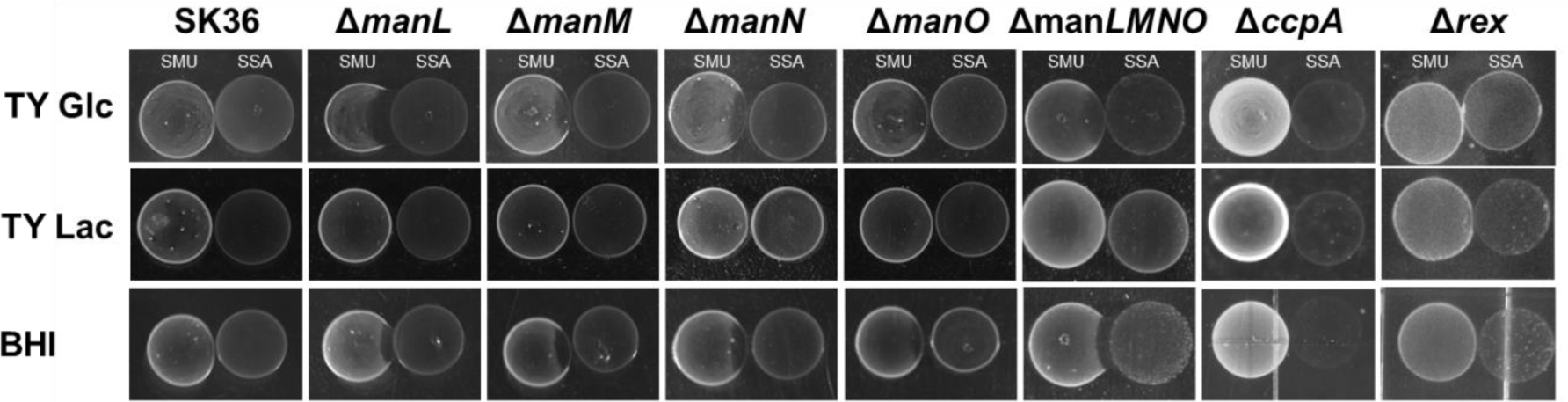
Antagonism of *S. mutans*. Cultures of SK36 and its mutant derivatives were each dropped onto the surface of BHI agar, or TY-agar plates supplemented with glucose or lactose and incubated for 24 hours in an aerobic atmosphere supplemented with 5% CO_2_. Cultures of *S. mutans* were then spotted to the left of the initial colonies and were incubated for an additional 24 hours. Each experiment was repeated three times, with a representative result being presented.

Pyruvate oxidase activity, by way of H_2_O_2_ excretion, has been shown to increase extracellular DNA (eDNA) release in *S. sanguinis* (13, 44). As eDNA is an important factor in biofilm development, we questioned the possibility of observed increases in H_2_O_2_ excretion altering the release of eDNA in these mutants. To evaluate this, supernatants from overnight cultures were mixed with a fluorescent DNA dye, SYTOX Green, and analyzed for relative eDNA content. Interestingly, multiple PTS deletion mutants with increased H_2_O_2_ excretion had less eDNA in their culture environment than the WT, with the most significant reduction being present in Δ*manL* (Fig 8A), a finding inconsistent with a previous report correlating H_2_O_2_ levels and eDNA release in the same bacterium (44). Notably, Δ*manM* and Δ*manO* produced WT levels of eDNA.

**Fig 8.**
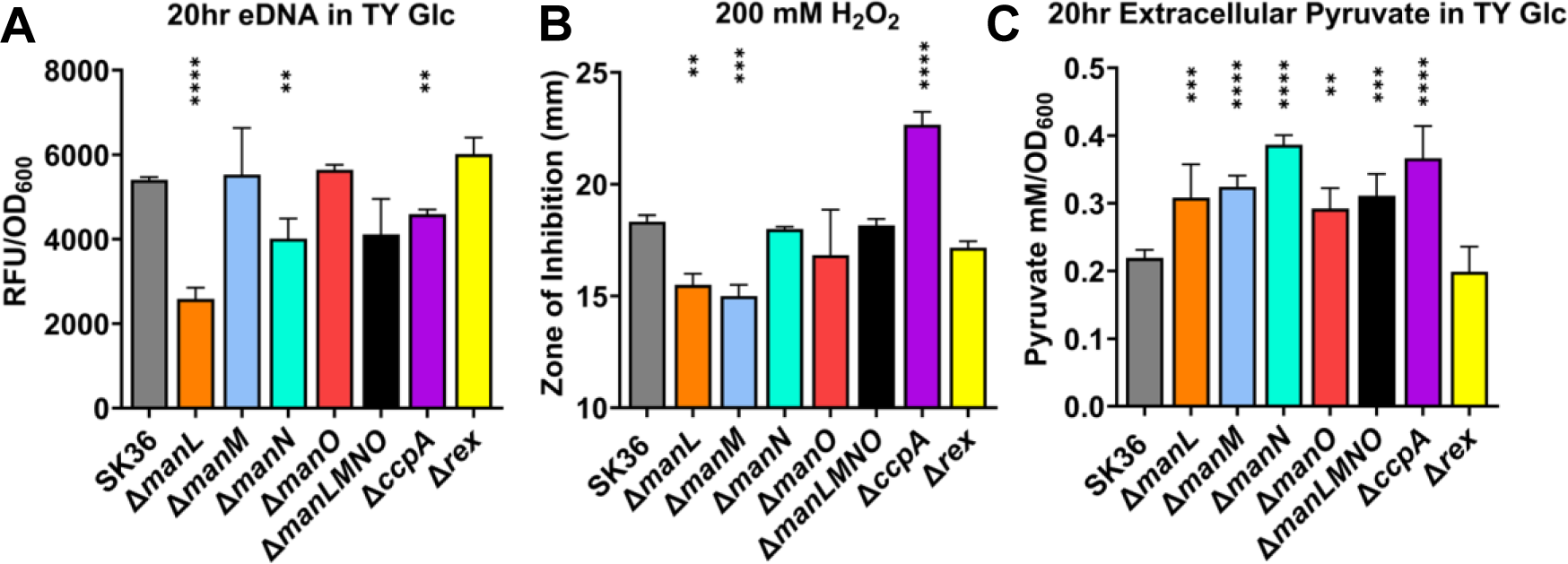
eDNA release (A), H_2_O_2_ stress survival (B), and pyruvate excretion (C) of SK36 and its mutant derivatives. To determine extracellular pyruvate and eDNA, cells were grown for 20 hours in TY medium supplemented with 20 mM glucose. Supernatants were then mixed with a fluorescent SYTOX Green stain and read at Ex485/Em528 to measure the relative amounts of extracellular DNA (A) or were used in an LDH-catalyzed reaction to measure pyruvate excretion (C). For determination of susceptibility to H_2_O_2_ stress, cells from mid-exponential phase were plated on BHI agar and H_2_O_2_ administered through disc diffusion (B). Results are the averages of three biological replicates, with error bars denoting standard deviations. Asterisks represent statistical significance compared to the wild type according to one-way ANOVA (*, *P* <0.05; **, *P* <0.01; ***, *P* <0.001; ****, *P* <0.0001).

As our previous study on Δ*manL* showed an upregulation of genes responsible for membrane biogenesis (32), we hypothesized this lack of eDNA release could at least in part be due to enhanced tolerance of oxidative stress. To test this theory, the EII^Man^ mutants were subjected to H_2_O_2_ stress through disc diffusion. Curiously, the observed phenotypes for EII^Man^ mutants were split into two groups, with strains Δ*manL* and Δ*manM* displaying increased tolerance, while Δ*manN*, Δ*manO*, and Δ*manLMNO* remained similar to the WT. Notably the Δ*ccpA* mutant was significantly more susceptible to H_2_O_2_ stress than all other strains (Fig 8B). As others have purported the possibility of extracellular pyruvate, derived from streptococcal metabolism, having a protective effect on H_2_O_2_ stress (40, 43), we examined extracellular pyruvate levels in similar culture conditions. When compared to SK36, all mutants tested displayed significantly increased pyruvate secretion levels in culture supernatants except for Δ*rex* (Fig 8C), but not in a manner correlating with observed eDNA release or H_2_O_2_ stress tolerance.

To test if this increased H_2_O_2_ excretion by EII^Man^ mutants contributed to their growth characteristics, SK36 and its mutant derivatives were grown in TY medium supplemented with glucose or fructose, a sugar not transported by the glucose-PTS, in the presence or absence of catalase to degrade H_2_O_2_ (Fig S8). With glucose being the main carbon source, addition of catalase significantly reduced the doubling time of all PTS mutants (excluding Δ*manO*), as well as improving the max OD_600_ in most (Table 1). This change was largely absent in fructose, a sugar that is not transported by the glucose-PTS. Therefore, these glucose-PTS mutants traded a reduction in PTS activity and growth rate for an enhancement in yield and competitive fitness, a strategy that could prove beneficial under carbohydrate-replete conditions.

**Table 1.** Growth characteristic of SK36 and its mutant derivatives. Presented are doubling time (T*_d_*, min, avg ± SD) and maximum OD_600_ (avg ± SD) after 24 hours of growth in TY media supplemented with 20 mM glucose or fructose, with and without 50 µg/ml catalase (cat) to degrade H_2_O_2_. Asterisks represent statistical significance when catalase is added, while pound symbols represent difference from the wild-type in the same condition, according to Welch’s *t*-test (*, *P* <0.05; **, *P* <0.01; ***, *P* <0.001; ****, *P* <0.0001; similar convention for ^#^).

| Strain | TY Glucose |  |  |  | TY Fructose |  |  |  |
| --- | --- | --- | --- | --- | --- | --- | --- | --- |
| | $T_d$ | | Max OD<sub>600</sub> | | $T_d$ | | Max OD<sub>600</sub> | |
|  | - cat | + cat | - cat | + cat | - cat | + cat | - cat | + cat |
| **SK36** | 89.06 ±<br>8.87 | 88.68 ±<br>6.16 | 0.60 ±<br>0.03 | 0.61 ±<br>0.04 | 65.85 ±<br>3.90 | 68.07 ±<br>7.49 | 0.63 ±<br>0.04 | 0.62 ±<br>0.04 |
| **$\Delta manL$** | 117.09 ±<br>2.29 ##### | 88.52 ±<br>5.66 **** | 0.85 ±<br>0.03 ##### | 0.94 ±<br>0.02 ****,<br>### | 71.23 ±<br>3.7 | 56.94 ±<br>4.47 ** | 0.66 ±<br>0.03 | 0.69 ±<br>0.05 |
| **$\Delta manM$** | 119.36 ±<br>5.88 ##### | 99.19 ±<br>7.36 **** | 0.89 ±<br>0.11 # | 0.91 ±<br>0.06 *, # | 83.19 ±<br>4.69 # | 87.43 ±<br>9.48 | 0.58 ±<br>0.08 | 0.58 ±<br>0.8 |
| **$\Delta manN$** | 128.93 ±<br>8.03 ##### | 95.82 ±<br>6.62 **** | 0.82 ±<br>0.05 ### | 0.81 ±<br>0.03 ****, ### | 75.95 ±<br>7.74 | 67.56 ±<br>3.86 | 0.62 ±<br>0.03 | 0.62 ±<br>0.03 |
| **$\Delta manO$** | 89.26 ±<br>6.25 | 92.48 ±<br>9.99 | 0.60 ±<br>0.07 | 0.59 ±<br>0.07 | 68.61 ±<br>5.41 | 62.06 ±<br>2.9 | 0.64 ±<br>0.04 | 0.62 ±<br>0.02 |
| **$\Delta manLMNO$** | 154.47 ±<br>5.72 ##### | 88.45 ±<br>8.69 **** | 0.83 ±<br>0.05 ### | 0.86 ±<br>0.06 ****, ### | 64.91 ±<br>6.76 | 57.17 ±<br>10.18 | 0.69 ±<br>0.03 | 0.64 ±<br>0.05 |
| **$\Delta rex$** | 85.22 ±<br>6.44 | 87.03 ±<br>4.88 | 0.61 ±<br>0.03 | 0.61 ±<br>0.02 | 87.13 ±<br>12.48 # | 44.85 ±<br>1.09 *, ## | 0.66 ±<br>0.02 | 0.73 ±<br>0.02 ***,<br>## |

### Deficiency in glucose-PTS alters bacterial pH homeostasis and the fate of pyruvate at multiple levels

Oral streptococci are important producers of both organic acids and ammonia, and CcpA and Rex have both been suggested to modulate pH homeostasis through multiple bacterial functions (18, 36, 37, 42). When deletion mutants of these regulators and the glucose-PTS were grown in TYG, EII^Man^ mutants (excluding Δ*manO*) generally displayed significantly increased pH in comparison to the WT, as did Δ*ccpA*, while Δ*rex* mirrored the WT (Fig 9A).

**Fig 9.**
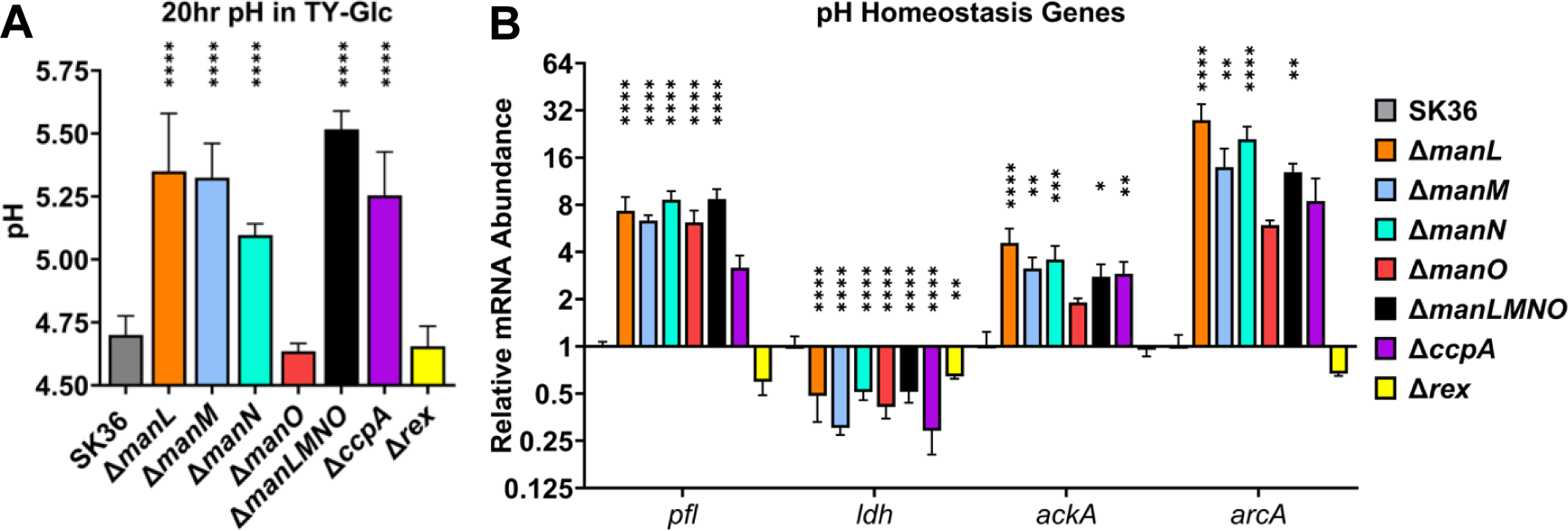
pH measurements (A) and transcription of metabolic genes (B). SK36 and its mutant derivatives were cultured in TY medium supplemented with 20 mM glucose for 20 hours for pH measurement (A) or to mid-exponential phase for RNA extraction (B). Results are the averages of three biological replicates, with error bars denoting standard deviations. Asterisks represent statistical significance compared to the wild type according to one-way ANOVA (*, *P* <0.05; **, *P* <0.01; ***, *P* <0.001; ****, *P* <0.0001).

To understand these phenotypes, a transcriptional profiling of genes relevant to the production of acids including formate (pyruvate formate lyase: *pfl*), lactate (lactate dehydrogenase: *ldh*), and acetate (acetate kinase: *ackA*), as well as ammonia (arginine deiminase: *arcA*), was conducted in these mutants (Fig 9B). Compared to the WT, expression of *ldh* was modestly decreased in all mutants, including Δ*manO* and Δ*rex*. Conversely, *pfl* and *ackA* were upregulated in all but Δ*rex*. Instead, Δ*rex* displayed slightly lower expression of both *ldh* and *pfl.* The *arcA* gene was also upregulated in all but Δ*rex*.

To corroborate this transcription analysis, three organic acids in spent media were measured. Concentrations of formate were increased by about 5-fold in PTS mutants (2-fold for Δ*manO*), matching the transcriptional data (Fig 10A). Similar increases in acetate levels were observed in the same mutants (Fig 10B). Despite the modest reduction in *ldh* transcripts in these PTS mutants, lactate levels were decreased by about 5-fold, except for Δ*manLMNO* which decreased by 2-fold, and Δ*manO* which remained unchanged (Fig 10C). Importantly, the acid profiles of Δ*ccpA* and Δ*rex* differed substantially from these PTS mutants: Δ*ccpA* produced WT levels of lactate, 3-fold higher acetate, and 2-fold lower formate; meanwhile, Δ*rex* displayed no discernible difference in acid output from the WT.

**Fig 10.**
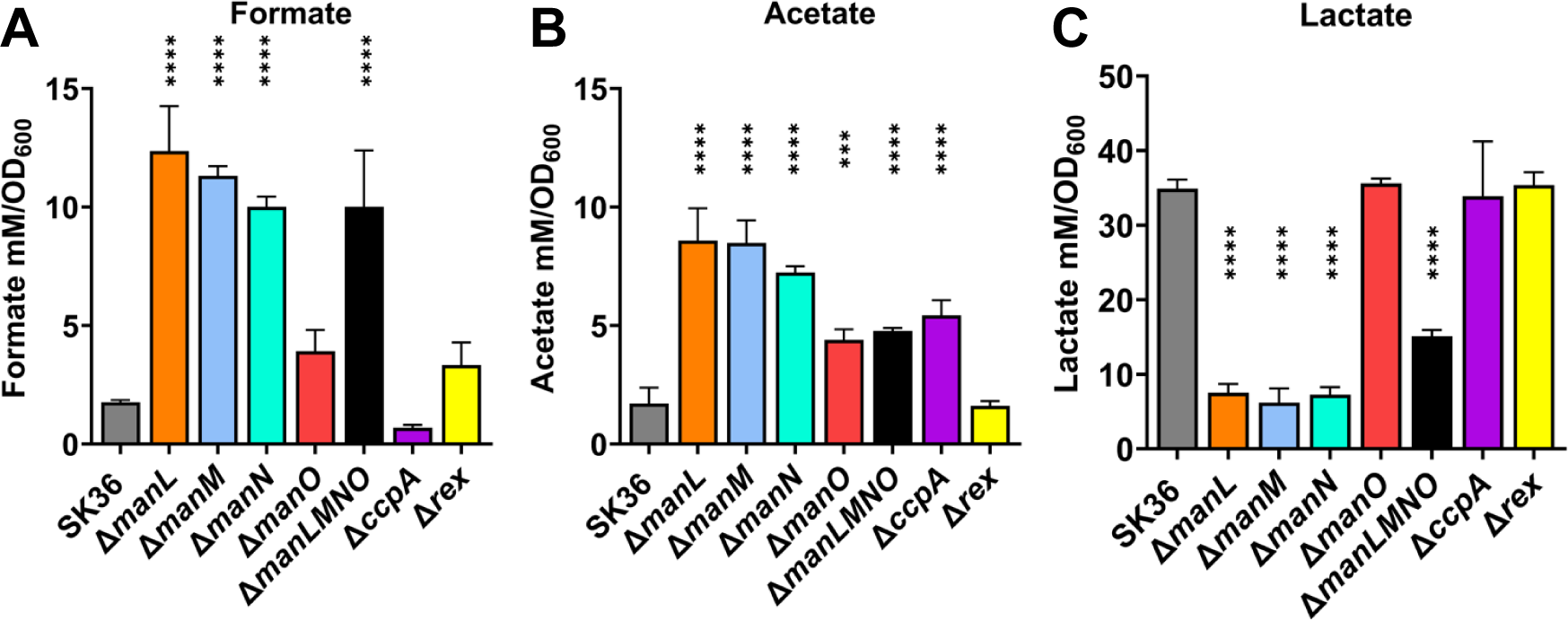
Organic acid excretion in SK36 and its mutant derivatives. Bacteria were grown to exponential phase in TV medium supplemented with 20 mM glucose. Supernatants were collected to measure levels of formate (A), acetate (B), or lactate (C). Asterisks represent statistical significance compared to the wild type according to one-way ANOVA (*, *P* <0.05; **, *P* <0.01; ***, *P* <0.001; ****, *P* <0.0001).

To ascertain the effects of ADS expression on bacterial pH homeostasis, SK36 and its mutant derivatives, *manL* and *manLMNO* were cultured in FMC with varying levels of arginine for 24 hours. As the arginine concentration decreased from the standard 400 µg/ml, the pH of the spent media also decreased gradually for both PTS mutants, a phenotype absent in the WT (Fig 11A); indicative of the role of arginine deiminase in moderating acidic pH. When comparing the growth yield over the arginine gradient however, both EII^Man^ mutants showed a reduced final OD_600_ when compared to the WT and the effect was relatively unchanged until arginine was completely absent in the media (Fig 11B), wherein the WT and both EII^Man^ mutants ceased to grow. As an *arcA* null mutant showed no such deficiency in the absence of arginine, we introduced an *arcA* deletion into the EII^Man^ mutants. In each case, deletion of *arcA* boosted the yield in FMC without arginine (Fig S9). It thus appears that the ADS activity may limit the amount of arginine that could otherwise be used for other crucial cellular processes, a scenario suggested previously (32).

**Fig 11.**
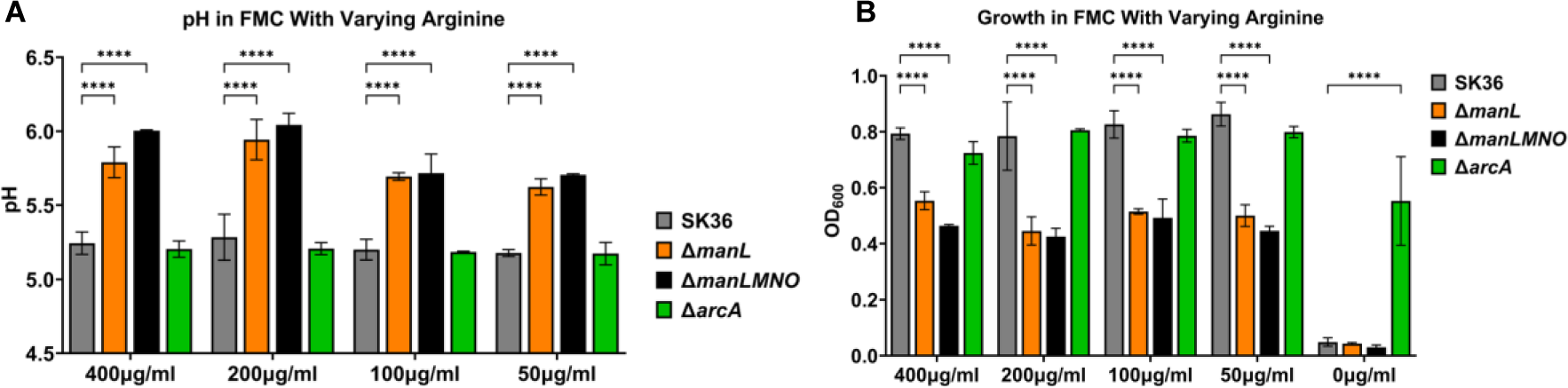
Resting pH (A) and final OD_600_ (B) of FMC cultures prepared with modified arginine levels. SK36 and its mutant derivatives were grown overnight in BHI medium and diluted 1:100 into FMC with 400, 200, 100, 50, or 0 µg/ml arginine, followed by 24 hours of incubation in an aerobic incubator maintained with 5% CO_2_. The 0 µg/ml condition is omitted in (A) due to lack of growth. Asterisks represent statistical significance compared to the wild type according to two-way ANOVA (*, *P* <0.05; **, *P* <0.01; ***, *P* <0.001; ****, *P* <0.0001).

### Deletion of HPr resulted in a marked reduction in cellular fitness

HPr, encoded by *ptsH*, is responsible for phosphorylating EIIA subunits and a cofactor for activating CcpA-mediated transcription regulation (45). In some related oral streptococci, such as *S. mutans*, a *ptsH*-null mutant is non-viable (30). To assess the role of HPr in the phenotypes of the EII^Man^ deletion mutants discussed thus far, we attempted to construct a deletion mutant of *ptsH* (Δ*ptsH*). The mutagenesis was carried out in both aerobic (with 5% CO_2_) and anaerobic environments, with each experiment generating substantial amounts of transformants. So far, only isolates from the aerobic condition contained the desired genetic change, whereas the isolates from the anaerobic condition appeared to harbor both the WT *ptsH* sequence and junctions indicative of an allelic exchange with the antibiotic cassette, as indicated by both PCR analysis (Fig S10) and WGS (Tables S4 and S5). WGS of one isolate from the aerobic condition revealed at least 6 SNPs (Table S5) and was selected for further analysis based on its growth phenotype. This Δ*ptsH* isolate showed greatly reduced growth capacity and tended to clump and fall out of suspension in liquid medium, which made it unsuitable to be assayed for any method normalized to growth state. When plated on TY-Glc PB or BHI PB agar, Δ*ptsH* produced very small amounts of H_2_O_2_ after 24 hours of incubation, with no clear bacterial colony structure. However, extended incubation of these plates showed this mutant capable of producing more H_2_O_2_ than the WT control (Fig 12).

**Fig 12.**
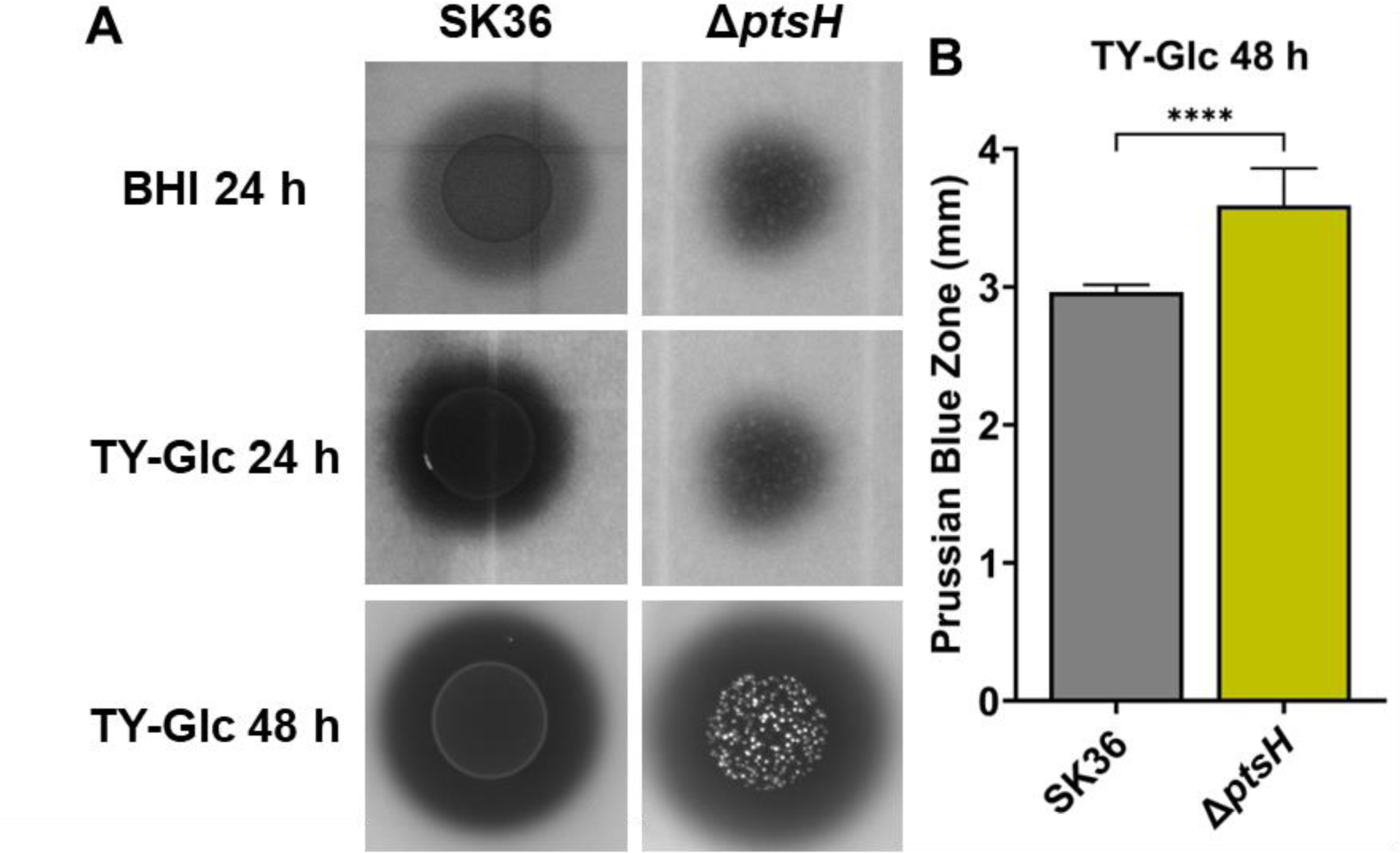
H_2_O_2_ production by SK36 and the *ptsH* mutant. Cultures of SK36 and its *ptsH* mutant derivative were each dropped onto BHI or TY-Glc agar plates containing PB reagent and incubated for either 24 or 48 hours in a 5% CO_2_ environment. All images were photographed under the same settings. Each experiment was repeated three times, with a representative result being presented (A). Each PB zone was measured from the edge of the bacterial colony to the edge of the PB precipitation at four locations using ImageJ software (B). Asterisks represent statistical significance compared to the wild type according to Student’s *t*-test (*, *P* < 0.05; **, *P* < 0.01; ***, *P* < 0.001; ****, *P* < 0.0001).

## Discussion

The ability of oral streptococci to colonize the oral surfaces is determined in large part by their ability to compete with other organisms and manage environmental stress, both of which are intimately tied to metabolism of carbohydrates. Multiple studies have demonstrated the complexity of carbohydrate-mediated metabolic regulation that works in concert with or independently of CcpA in streptococcal pathogens (20, 30, 46–48). What is not well understood is whether a similar function of the PTS exists in a group of the most abundant species in the oral biofilm, commensal streptococci, and its influence in microbial homeostasis. Built on our previous research revealing the phenotypic and transcriptomic changes when EIIAB^Man^ (*manL*) was deleted (32, 34), here, we conducted a systematic analysis on the contribution of individual components of the glucose-PTS to physiology and fitness.

This study further substantiated some of our previous findings in physiology and fitness by demonstrating that mutations in any of the 4 main subunits of EIIABCD^Man^ reprogrammed the central carbon metabolism that increased products that are important to antagonism (H_2_O_2_), membrane biogenesis (acetyl-CoA and glycerol metabolism), bioenergetics (ATP), and pH homeostasis (ammonia) (32). The culmination of these metabolic adjustments led to profound alterations in cellular physiology. These mutants produced enhanced yield (max OD_600_) and superior fitness when competing against major oral pathobiont *S. mutans*. Carbon flux through the pyruvate node appeared to have shifted from a mainly homolactic strategy to a mixed acid fermentation strategy, paired with increased arginine deiminase activity that excreted pH-buffering ammonia. This type of metabolic shift is often considered an indication of alleviation in CcpA-mediated CCR and an outcome of decreased carbon flux (16, 19, 21). On the contrary, our work demonstrated the critical distinctions between PTS-mediated catabolic regulation and what is effected by CcpA, or Rex, as CcpA-mediated CCR is insufficient to explain these novel phenotypes (16, 32). While Δ*rex* failed to show any significant phenotype in most of our assays, mutations in the PTS components resulted in a range of phenotypes in growth, acid byproducts, and antagonism that were not only different from that induced by deletion of *ccpA* but, more importantly, included variations such as increased PTS activity and relieved CCR in ManNA91E strains, which pointed to a direct involvement of the PTS transporter in the underlying mechanism.

Nonetheless, the phenotypic similarity among these EII^Man^ deletion mutants studied thus far may imply a certain mechanism that transduces the signal of the carbon flow into regulation. One possibility is HPr which has been implicated in different modes of gene regulation independently of CcpA (30), and another is EIIAB^Man^ (ManL) which potentially functions in response to the phosphorylation status of the PTS (49). To further test the involvement of ManL, SNP mutations were introduced to block the phosphorylation of subunits A and B, separately, by switching the phosphorylating residues His13 of EIIA^Man^ to alanine, and His185 of EIIB^Man^ to alanine. These strains however showed no discernible difference from the complete *manL* deletion mutant in terms of H_2_O_2_ excretion, pH, and lactate excretion (Fig S11).

We were successful in constructing a deletion mutant of HPr (Δ*ptsH*) in *S. sanguinis,* although it is likely that compensatory mutations are needed to keep it viable under the lab conditions. In agreement with its function as the key member of the PTS, this mutant of HPr was highly debilitated in growth. Perhaps it is not surprising that a mutant that is critically impaired in sugar transport should produce less H_2_O_2_ simply for lack of substrate for the pyruvate oxidase SpxB. However, the fact that Δ*ptsH* eventually produced more H_2_O_2_ than the WT, after two days on agar plates, does provide some partial support to the theory that HPr is involved in regulating these fitness phenotypes affected by mutations of the EII^Man^ operon. Past genetic research has indicated that *ptsH* may not be an essential gene for the species of *S. sanguinis* (50, 51). Due to its poor growth however, we were not able to reliably compare Δ*ptsH* with the rest of the PTS mutants in several other metabolic assays. It is also worth noting that repeated attempts under anaerobic conditions failed to generate a similar *ptsH* mutant. Considering the fact that H_2_O_2_, as a product of oxygen metabolism, can act as a mutagen and increase the rate of spontaneous mutation, this absence of compensatory mutations under anaerobic conditions could simply be due to the lack of H_2_O_2_. Interestingly, several of these isolates (3 out of 4 sequenced) from anaerobic conditions did contain an SNP within the *clpX* gene, which is responsible for negatively regulating oxidative stress genes (52, 53). Further research is needed to determine the significance of these compensatory SNPs in relation to HPr-mediated functions in both aerobic and anaerobic conditions. Related to this subject are the spontaneous mutations identified in the ManNA91E mutants. While it is highly likely that the observations made of these mutants are largely due to the intended mutagenesis of the target gene, we cannot rule out the possible contributions by other present SNPs (Table S2), which can be addressed in future studies.

PTS-mediated CCR and CCR controlled by CcpA are intertwined as alterations in carbon flux sensed by the PTS can translate into changes in metabolic intermediates such as FBP which can trigger Ser-phosphorylation of HPr and activate the function of CcpA. This provides an explanation for some of the overlapping phenotypes between these two systems observed in this study, e.g., expression of *spxB* and *arcA*. A key distinction between these two mechanisms, however, is illustrated by a previous finding in *S. mutans* that while PTS responds to glucose in the μM to low mM range, CcpA requires at least 3-5 mM of monosaccharides to trigger its function (26). We have previously demonstrated the involvement of both mechanisms in the regulation of catabolic genes in *S. mutans* (26) as well as in *S. gordonii* (27). Based on the acid profile (Fig 10), H_2_O_2_-mediated antagonism (Fig 7), and stress tolerance (Fig 8) between Δ*ccpA* and the PTS mutants, it is likely that a similar, PTS-mediated, CcpA-independent regulation is governing the key functions responsible for both competition and stress tolerance in *S. sanguinis*. With glucose being present in saliva at steady-state levels (∼80 μM, notably higher in hyperglycemic individuals) presumably sufficient to trigger PTS-mediated CCR even during “famine” period (26, 54), these novel findings in streptococcal metabolic regulation could prove significant to our effort to modulate the microbial homeostasis in the oral cavity. We also have reason to believe that PTS has a similar influence on the acid profile of *S. mutans* (32). We speculate that a suitable small molecule, selected based on its ability to interact with and influence the function of the PTS transporter, may be applied to the oral microbiome for improving acid profile, pH homeostasis, and the overall robustness of the commensals. Furthermore, with arginine being researched as a highly promising prebiotic candidate against dental caries, the observation that arginine deiminase activity in these bacteria reduced their fitness in an arginine deplete condition warrants further study to assess the significance of salivary arginine levels in the health status of the oral microbiome.

Last, we remain open to the possibility that some of these PTS-dependent phenotypes were the outcomes of post-transcription regulation, including allosteric regulations mediated by protein: protein and protein: metabolite interactions. In particular, the gene *manO* could prove to be of unique significance in regulation, in part because deleting it did not result in drastic phenotypes indicative of a structural role in the function of the PTS. Similar accessory genes encoded as part of the PTS operon have been reported previously (55). It is possible that specific conditions are yet to be identified under which the full function of ManO can be revealed.

Overall, these findings in fitness phenotypes are indicative of glucose-PTS playing a direct and much larger role in regulating oral streptococcal pathophysiology in the plaque environment than previously thought. It is probable that similar regulations and phenotypic outcomes are conserved in other oral streptococci or non-streptococcal lactic acid bacteria. We interpret these results to underscore the necessity of further study into not only the impact of glucose-PTS on regulations at transcriptional and enzymatic levels, but also into understanding these effects at a multi-species level. This influence on bacterial fitness could allow us to predict and prevent caries by acting on both commensal and pathogenic sides of the community homeostasis.

## Materials and Methods

### Bacterial strains and culture conditions

Strains (Table 2) including SK36 of *S. sanguinis,* its mutant derivatives, and UA159 of *S. mutans* were maintained on BHI (Difco Laboratories, Detroit, MI) agar supplemented with 50 mM potassium phosphate buffer, pH 7.2. The antibiotics erythromycin (Em; 10 µg/ml), and kanamycin (Km; 1 mg/ml) were used in agar plates to select for antibiotic-resistant transformants when necessary. BHI liquid medium was routinely used for the preparation of batch starter cultures, which were then diluted into BHI, tryptone-yeast extract (TY; 3% tryptone and 0.5% yeast extract), or the chemically defined medium FMC modified to contain different carbohydrates at specific concentrations. A tryptone-vitamin (TV) medium was used for biochemical assays to avoid contaminating metabolites from yeast extract. Liquid cultures were harvested at specified growth phases by centrifugation (Sorvall Legend XTR) at 3800 x *g* at 4°C for 10 minutes. The cells or supernatants were used immediately for biochemical experimentation or stored at -80°C. To study growth characteristics, bacterial starter cultures were diluted into TY medium containing various carbohydrates and loaded onto a Bioscreen C system, where wells were each overlaid with 70 µl mineral oil, and cultures were maintained at 37°C.

**Table 2.**
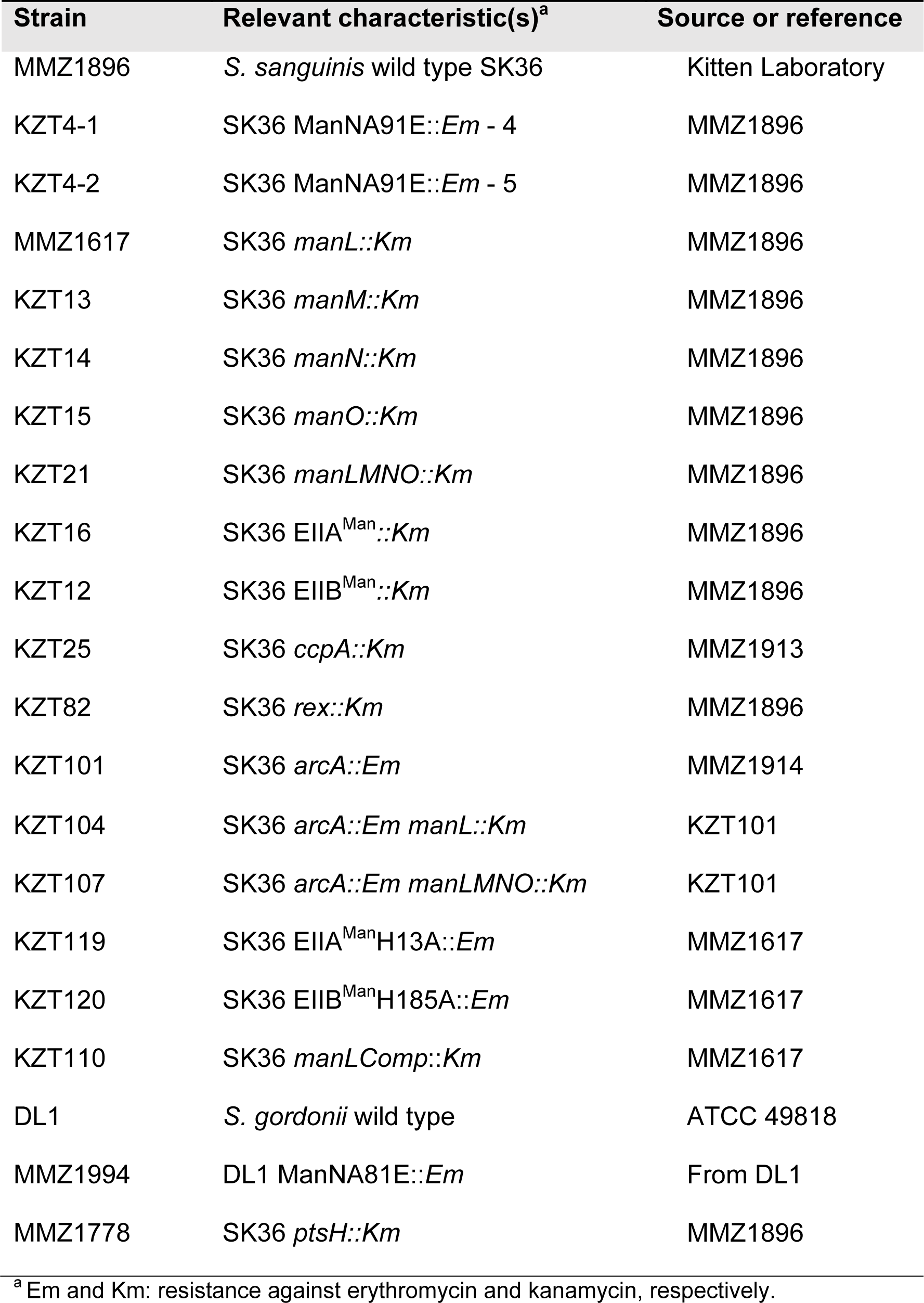
Strains used in this study.

To observe the role of arginine on cellular growth, bacterial starter cultures were diluted into FMC medium containing various concentrations of arginine and grown for 24 hours. OD_600_ and resting pH were measured in the resulting cultures.

### Construction of SNP and deletion mutants

Standard molecular cloning protocols were followed in the manipulation of various forms of DNA products, and transformation assays were performed primarily according to an established procedure that utilizes the natural competence phenotype displayed by *S. sanguinis* (34). All DNA oligonucleotides were synthesized by Integrated DNA Technologies (Coralville, IA) and are listed in Table S6. To construct the ManNA91E mutant, a 1-kbp DNA fragment was generated using a PCR product originating from the passaged strain containing the SNP in question. Subsequently, this 1 kbp mutator DNA was used together with a helper plasmid, p*pfl*::*gfp*::*erm*, in a co-transformation assay to introduce into the SK36 background the ManNA91E mutation and resistance to erythromycin. The resulting mutants were screened by allele-specific PCR (Table S6) (56). Chromosomal DNA was extracted from bacterial cells using a Wizard Genomic DNA purification kit (Promega, Madison, WI) and submitted to the SEQCENTER (Pittsburgh, PA) for WGS (Illumina) analysis and variant calling (Tables S1, S2, S4, and S5). All genomic DNA sequences received have been deposited in the Sequence Read Archive (SRA) under the Bioproject code PRJNA1133684 and assigned accession numbers SRX25252496, SRX25252497, SRX25252498, SRX25252499, SRX25252500, SRX25252501, and SRX25252502.

Deletion mutants were constructed using a modified allelic exchange strategy (57). Briefly, two DNA fragments flanking the desired region of deletion were generated by PCR and included ends overlapping a kanamycin or erythromycin resistance cassette. These two flanking DNA fragments and one antibiotic cassette were then incubated together in a Gibson assembly reaction for isothermal ligation (New England BioLabs, Beverly, MA), with the cassette replacing the target gene. This ligation product was then introduced into early exponential phase cells and incubated for 3 hours to mutate the genome through homologous recombination. The resulting mutants were selected on agar plates containing corresponding antibiotics and verified through PCR and Sanger sequencing, or WGS if necessary.

### PTS assay

The capacity of the bacterium to transport and phosphorylate carbohydrates was assessed using the PTS assay as previously described (47, 58). Briefly, bacterial cultures were permeabilized and their ability to phosphorylate specific carbohydrates in a PEP-dependent fashion, which releases pyruvate, was coupled with the reduction of pyruvate at the expense of NADH. To avoid the effects of endogenous NADH oxidase (*nox*) oxidizing the substrate of the assay, a *nox* deletion was introduced to the backgrounds of both SK36 and ManNA91E and the resultant strains were used in this assay.

### RNA extraction and qPCR

Bacterial cultures (7 ml) from the mid-exponential phase (OD_600_ = 0.5 to 0.6) were harvested and treated with RNAprotect reagent (Qiagen, Germantown, MD), and the cell pellet, if not immediately processed, was stored at -20°C. Bacterial cells were resuspended in 300 µl of 50/10 TE (Tris-EDTA 50/10 mM) with 0.4% SDS, together with an equal volume of acidic phenol and a similar volume of glass beads, and disrupted by bead-beating for 1 min at 4°C. After 10 min of centrifugation (Labnet Prism Microcentrifuge) at 18,000 x *g* at room temperature, the clarified aqueous layer was removed and processed using an RNeasy minikit (Qiagen) for extraction of total RNA. While loaded on the membrane of the centrifugal column, the RNA sample was treated with RNase-free DNase I solution (Qiagen), twice, to remove genomic DNA contamination. To synthesize cDNA, 1 µg of each RNA sample was used in a 20-µl reverse transcription reaction set up using the iScript Select cDNA synthesis kit (Bio-Rad), together with gene-specific reverse primers (Table S6) used at 200 nM each. Primer for the housekeeping gene *gyrA* was used as an internal control in all cases (59). After a 10-fold dilution with water, the cDNA was used as a template in a quantitative PCR (qPCR) prepared using an SsoAdvanced Universal SYBR Green Supermix and cycled on a CFX96 real-time PCR detection system (Bio-Rad), following the supplier’s instructions. Each strain was represented by three biological replicates, and each cDNA sample was assayed at least twice in the qPCR. The relative abundance of each mRNA was calculated against the housekeeping gene using a ΔΔCq method (60).

### H_2_O_2_ measurement and plate-based competition assay

The relative capacity of each strain to produce H_2_O_2_ was assessed using indicator agar plates on the basis of Prussian Blue (PB) formation (61). Briefly, a TY agar (1.5%) base was prepared with the addition of FeCl_3_·6H_2_O (0.1%) and potassium hexacyanoferrate (III) (0.1%). After autoclaving, various carbohydrates were added at specified amounts before pouring. Each strain was cultivated overnight in BHI, washed twice with sterile PBS, and a volume of 10 μl was pipetted onto the agar surface and then incubated for 20 hours to allow bacterial growth and development of PB precipitation. The PB zones were measured from colony edge to zone edge using ImageJ Fiji (62). Each strain was tested at least three times on 2 separate plates containing a fixed volume of agar medium for optimal comparison.

H_2_O_2_ excretion in the culture supernatant was quantified using a liquid quantification (63). Briefly, bacterial cultures (3.5 ml) from early exponential phase (OD_600_ = 0.3 to 0.4) were placed in an orbital shaker at 250 RPM at 37°C for 30 minutes to allow for thorough aeration. Cultures were spun down (Sorvall Legend XTR) at 3800 x *g* at 4°C for 5 minutes. 650 µl of supernatant was incubated with 600 µl of reaction buffer (2.6% a-amino-antipyrine, 1.6% saturated phenol in H_2_O) for four minutes at RT. 3.25 µl of 5 U/ml horseradish peroxidase was added and incubated at RT for 20 minutes. A H_2_O_2_ standard in the range of 0.068 to 0.407 mM was prepared freshly in TY base medium. The light absorbance of mixtures at OD_510_ was then read and normalized to the optical density (OD_600_) of the resulting culture.

Plate-based antagonism assays (64) were carried out to test the interactions between *S. sanguinis* and *S. mutans* UA159 on various carbohydrate sources using TY-agar supplemented with either 20 mM glucose or lactose, or BHI-agar. Overnight cultures of *S. sanguinis* strains were dropped onto the agar first and then incubated for 24 hours at 37°C in a 5%-CO_2_ aerobic incubator, followed by spotting of *S. mutans* UA159 near the *S. sanguinis* colony. Plates were incubated for another 20 hours before photographing. Each interaction was tested at least three times. As a negative control, catalase (Sigma-Aldrich) was used at 10 mg/ml and spotted on the media before *S. mutans* culture to inactivate any H_2_O_2_ excreted by *S. sanguinis*.

### Survival under exogenous H_2_O_2_ stress

To measure strain viability in an oxidative stress environment, mid-exponential phase cultures were plated as a lawn on BHI agar plates and allowed to dry. Autoclaved paper discs were then placed on top of the media and 20 µl of 200 mM H_2_O_2_ was pipetted onto the discs and allowed to dry. Plates were then incubated in 5% CO_2_ at 37°C. After 24 hours, plates were photographed and the diameter of the resulting zones of inhibition were measured using ImageJ Fiji (62). Each strain was tested at least three times on 2 separate plates containing a fixed volume of agar medium for optimal comparison.

### Biochemical assays for measurement of metabolites

Overnight cultures of SK36 and respective mutants in BHI were diluted 20-fold in TV containing 20 mM glucose and subcultured until OD_600_ reached 0.5. Immediately after centrifugation (Sorvall Legend Micro 21R) at 4°C at 10,000× g for 5 min, cell supernatants were extracted and frozen at -20°C until analysis. Concentration of Lactate was measured using an established lactate assay protocol utilizing LDH activity (65). Concentrations of acetate and formate were measured using an acetate assay kit (K-ACETAK, Neogen, Lansing, MI), and a formate assay kit (EFOR-100, BioAssay Systems, Hayward, CA) respectively, following manufacturers’ protocols. To avoid contaminating metabolites from yeast extract, all cultures were prepared using TV-glucose medium. Each biochemical assay was conducted using at least three biological replicates, alongside a standard prepared using known concentrations of the substrate of interest.

To measure pyruvate levels, we used an LDH-catalyzed reaction that coupled the reduction of pyruvate with the oxidation of NADH by monitoring the optical density at 340 nm (OD_340_). The assay was performed by mixing 10 µl of culture supernatant and 90 µl of enzyme solution, which included 10 units/ml of lactate dehydrogenase (Sigma) and 100 mM NADH in a 100 mM sodium-potassium phosphate buffer (pH 7.2) supplemented with 5 mM MgCl_2_. The reaction mixture was incubated at room temperature for 15 min before spectrometry, with the light source set at UV range (OD_340_). To rule out the influence on the assay by background NADH leakage from the cell, a control without the addition of LDH or NADH was included for each sample. A sodium pyruvate standard in the range of 0.2 to 1.0 mM was prepared freshly in TY base medium. The final measurements of pyruvate concentration were normalized against the optical density (OD_600_) of each culture.

### Statistics

Statistical analysis of data was carried out using the software of Prism (GraphPad of Dotmatics, San Diego, CA).

## Acknowledgments

This work was supported by a grant DE12236 from NIDCR and a startup fund to LZ from University of Florida. ZAT was supported in part by a T90 training grant DE021990 from NIDCR. We acknowledge Payam Noeparvar for technical assistance in performing various assay preparations. LZ conceptualized and supervised the study; ZAT and DNP performed the experiments; ZAT analyzed the data; ZAT and LZ wrote the manuscript.

